# Optogenetic control of lysosome function

**DOI:** 10.1101/2023.08.02.551716

**Authors:** Nikolay S. Ilyinsky, Sergey M. Bukhalovich, Diana F. Bagaeva, Alexey A. Alekseev, Semen V. Nesterov, Fedor M. Tsybrov, Andrey O. Bogorodskiy, Sofia F. Nazarova, Vadim A. Alekhin, Olga V. Moiseeva, Anastasiya D. Vlasova, Kirill V. Kovalev, Anatoliy E. Mikhailov, Andrey V. Rogachev, Ernst Bamberg, Valentin I. Borshchevskiy, Valentin Gordeliy

## Abstract

Lysosome protective, metabolic, signaling functions are highly dependent on their pH. A lack of tools of high spatial and temporal resolution for pH control is a bottleneck of lysosome related cell research. Light-driven inward proton pump *Ns*XeR, targeted to the lysosomes of mammalian cells, produces lysosome alkalization simply by light. Complementary use of outward proton pumping Arch3 rhodopsins in lysosomes offers an approach to vary pH in a range from around 5 to 6.5 in both directions (de-acidification and acidification). Lyso-*Ns*XeR optogenetics efficiency was demonstrated, in particular, by its ability to inhibit lysosome proteolytic enzymes. Thus, optogenetic monitoring and regulation of the lysosome function, through pH control over a wide range, could serve as an approach to studying fundamental cell processes, and rational drug design.

## Introduction

Lysosomes are indispensable organelles of the cells serving as a waste disposal/food digesting system cleaving used cellular components, macromolecules, and viruses through autophagy and endocytosis. They are involved not only in material and energy metabolism but also in other organelles regulation, plasma membrane repair, programmed cell death, secretion and signaling (Ballabio and Bonifacino, 2020). In general, pH of cellular compartments is a vital physiological parameter of homeostasis virtually in all cellular compartments, and pH alterations are associated with severe diseases (Weisz, 2003). Acidic environment is especially crucial for endocytic/phagocytic and autophagic pathways organelles, lysosomes in particular. Highly acidic environment (pH 4.5 – 5.0) in lysosomal lumen is required for various purposes like the efficient operation of hydrolytic enzymes (that degrade damaged biomolecules or nutrients), iron uptake, receptors recycling, lysosome positioning and release the viral genome. Low pH is also necessary for some amino acid transporters participating in the export of digested nutrients and metabolic signaling.

The ion balance in lysosomes is maintained by chloride, calcium, sodium, other ion transporters and by a proton pumping vATPase (Riederer et al., 2023), which plays a major role in the pH regulation. Inhibition of vATPase, lysosome membrane permeabilization, garbage accumulation lead to deacidification of lysosomes, which, in its turn, has severe consequences for human health. Dysregulations of lysosomes lead to impairment of energetics, proteostasis, and some other cell functions causing severe diseases, including cancer, neurodegeneration, age-related macular degeneration, infectious diseases, and metabolic disorders (Bonam et al., 2019). On the contrary, stabilization of lysosome activity, first of all acidic pH, is important for the embryo development and longevity (Adam Bohnert and Kenyon, 2017; Leeman et al., 2018; Sun et al., 2020). On the other hand, controlled lysosome alkalization could be used as simulation of diseases for the studies of their pathogenesis process. Many physiological cellular responses induced by lysosome alkalization (like lysosomal-mitochondrial calcium exchange, lysosome-to-nucleus signaling via TFEB, ER remodeling, metabolic adaptation and rewiring, and virus egress) should be re-investigated under spatio-temporally precise lysosome pH control. Lysosome alkalization could prevent viruses infectivity, cancer invasiveness and chemotherapy resistance, systemic autoimmune diseases. The analysis of lysosome pH influence on cellular processes carried out in the Supplementary Note 1.

Thus, manipulation of lysosomal pH is a key for the studies of their function and also therapeutic intervention. Optogenetic deacidification of lysosomes would be a missing, problem-solving approach to the studies of lysosome functions and dysfunctions. Optogenetics is already widely applied for the multimodal remote control of neurons by use of microbial rhodopsins (Gordeliy, 2022), ion pumps, and ion channels selectively expressed in the plasma membrane of the cells (Adamantidis et al., 2015). Though the rhodopsin optogenetic approach in cell organelles is a minimally invasive treatment, few, comparing to neuroscience applications, examples of intracellular optogenetic tools exist. There are rhodopsin-assisted studies of synaptic vesicles and lysosomes (Rost et al., 2015), endosomes (Kakegawa et al., 2018), mitochondria (Berry and Wojtovich, 2020), and endoplasmatic reticulum (Asano et al., 2018). For example, functional incorporation of rhodopsins to the cell organelles allows optogenetic control of the electrochemical gradient by channelrhodopsin in mitochondria (Tkatch et al., 2017). Also, application of the outwardly directed proton pump Arch3 in synaptic vesicles enables light control of vesicular acidification and neurotransmitter accumulation (Rost et al., 2015). Recently optogenetic approaches to cytosol alkalization and acidification were developed (Donahue et al., 2021; Nakao et al., 2022; Vlasova et al., 2023). The reason for limited representation of optogenetics in the control of different cellular compartments is unresolved difficulties of specific targeted expression of optogenetic tools (Garita-Hernandez et al., 2018). Thus, specific organelle optogenetics is a promising emerging area.

In our work, we expand the optogenetics toolbox to control lysosome functions not only through their acidification but also through their alkalization via expression of outward and inward light-driven pumps, correspondingly, in lysosomes of mammalian cells. We demonstrate the efficiency of lysosome optogenetics with an inward proton pump, xenorhodopsin from the nanohalosarchaeon *Nanosalina* (*Ns*XeR) (Shevchenko et al., 2017), enabling alkalization of the organelle up to 1 pH unit. It makes it possible to control lysosome functions through optogenetic regulation of pH in wide physiological and pathological ranges (Leung et al., 2019), confirmed here by controlled inhibition of lysosomal pH-dependent enzyme Cathepsin B. Optogenetic control of pH in a wide range (from acidic to alkaline) opens a new way to the studies of key processes in which lysosomes are involved, particularly those driven by alkalization of lysosomes. Since acidification is required for the function of different cellular organelles (Weisz, 2003), the optogenetics introduced here with light-driven inward proton pumps might have applications beyond lysosomes, for the studies of cellular functions and dysfunctions dependent on the pH state of other organelles.

## Results

### Lyso-Arch3, established optogenetic tool for lysosomal acidification: determination of the corresponding pH change in absolute units

There is an optogenetic approach of lysosome acidification by established previously lyso-pHoenix construct (named here as lyso-Arch3), containing an outward proton pump archaerhodopsin from Halorubrum sodomense Arch3, which was shown to localize in the lysosome membrane and could induce transient lysosome acidification in Bafilomycin A1 (BafA1) treated cells (Rost et al., 2015).

To understand the real performance of this optogenetic approach and be able to rationally plan future optogenetic studies of lysosome and lysosome-related processes in the cells, one should know the pH shift of lyso-Arch3 in absolute pH units. We expressed the outward rhodopsin proton pump Arch3 in lysosomes of the HEK293T cells (see experimental pipeline on Figure 1A), as described by Rost *et al*. (Rost et al., 2015). Cells were incubated with Bafilomycin A1 (BafA1) overnight before the experiment to achieve synchronized alkalization of lysosomes (Figure 1B), that was revealed by yellow lysosomes having high fluorescence of pHluorin (pH-sensitive green fluorescent protein in lyso-Arch3 construct), mixing with mKate2 (pH-insensitive red fluorescent protein in lyso-Arch3 construct) signal.

**Figure 1.**
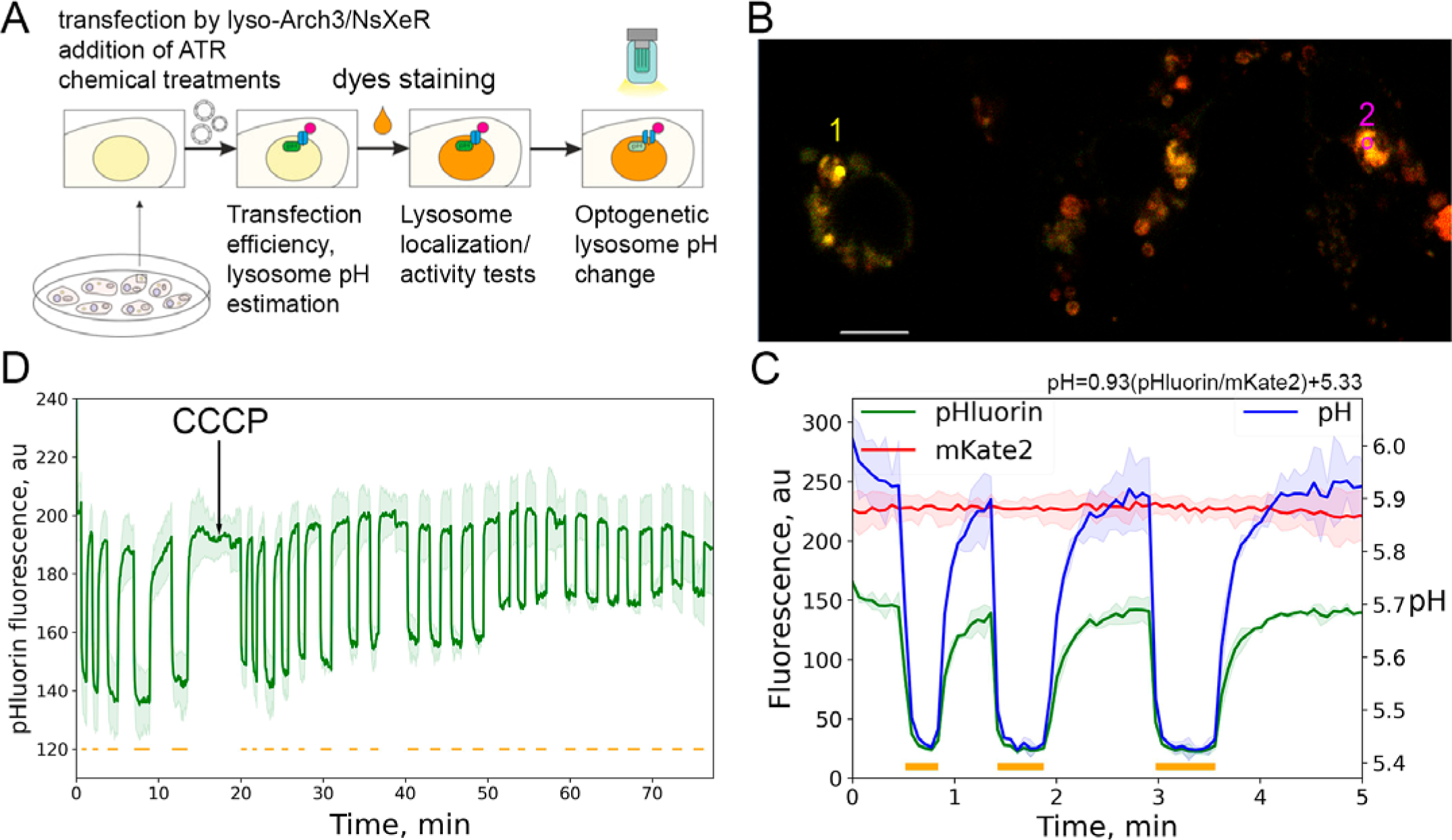
Lyso-Arch3 application in HEK293T cells treated with vATPase inhibitor BafA1. (A) Experimental pipeline of fluorescent detection and optogenetic manipulation of lysosomal pH, activity. (B) Image of confocal fluorescence microscopy showing co-expression of pH-sensitive green fluorescent protein pHluorin and pH-insensitive red fluorescent protein mKate2, with high green (as result yellow) fluorescence in BafA1 de-acidified lysosomes. Scale bar 10μm. (C) Demonstration of lysosomal acidification (by pHluorin sensor, green curve) under illumination. Fluorescence is measured at regions 1 and 2 highlighted on (B). Time intervals of illumination by 590nm light are highlighted by orange lines. The absence of mKate2 fluorescence change (red curve) proves that lysosomes are not moving under illumination. The fluorescence signals were corrected on background to remove backlight from LED. Blue curve represents maximum lysosomal ΔpH (calculated by calibration) under described conditions. (D) The level of lysosomal acidification by lyso-Arch3 decreases during the incubation with the protonophore 50uM CCCP. All experiments with lyso-Arch3 were done after overnight treatment with 50nM of BafA1. Time curves presented as mean (for the data from five cells)±standard deviation (SD).

First, we proved that the pHluorin fluorescence change (Figure 1, green curve) is due to the change of pH rather than the lysosomal movement (Figure 1, no change in red curve for mKate2, see also Supplementary Movie 1).

Second, by pHluorin calibration (Sankaranarayanan et al., 2000) it was shown that Lyso-Arch3 can decrease pH in BafA1-treated HEK293T by about 0.5 units (Figure 1C, blue curve). We should note, that precise determination of the maximum drop of pH is limited since super-ecliptic pHluorin loses linearity of the pH-response below pH ∼5.5 and is insensitive below pH ∼5.0 (Sankaranarayanan et al., 2000).

Third, the experiments showed that the emerging ΔpH produced by lyso-Arch3 disappears under the protonophore treatment (Figure 1D), proving that proton ions pumping by lyso-Arch3 is responsible for pHluorin fluorescence changes rather that over ions or artifacts.

### Lyso-*Ns*XeR: new optogenetic tool for the pH increase in lysosomes

The main goal of our work was to develop an optogenetic tool for reversible increase of pH of lysosomes. For this end, we expressed an inward proton pump *Ns*XeR (Shevchenko et al., 2017) in a fusion construct CD63N-se-pHluorin-*Ns*XeR-mKate2-βHK-CD63C (hereafter lyso-*Ns*XeR, Figure 2A) with the pH sensitive green fluorescent protein super-ecliptic (se-) pHluorin (hereafter pHluorin, in lysosome lumen), and the pH insensitive red fluorescent protein mKate2 (facing the cytosol). This approach has been used previously with lyso-Arch3 construct, described in previous section (Rost et al., 2015).

**Figure 2.**
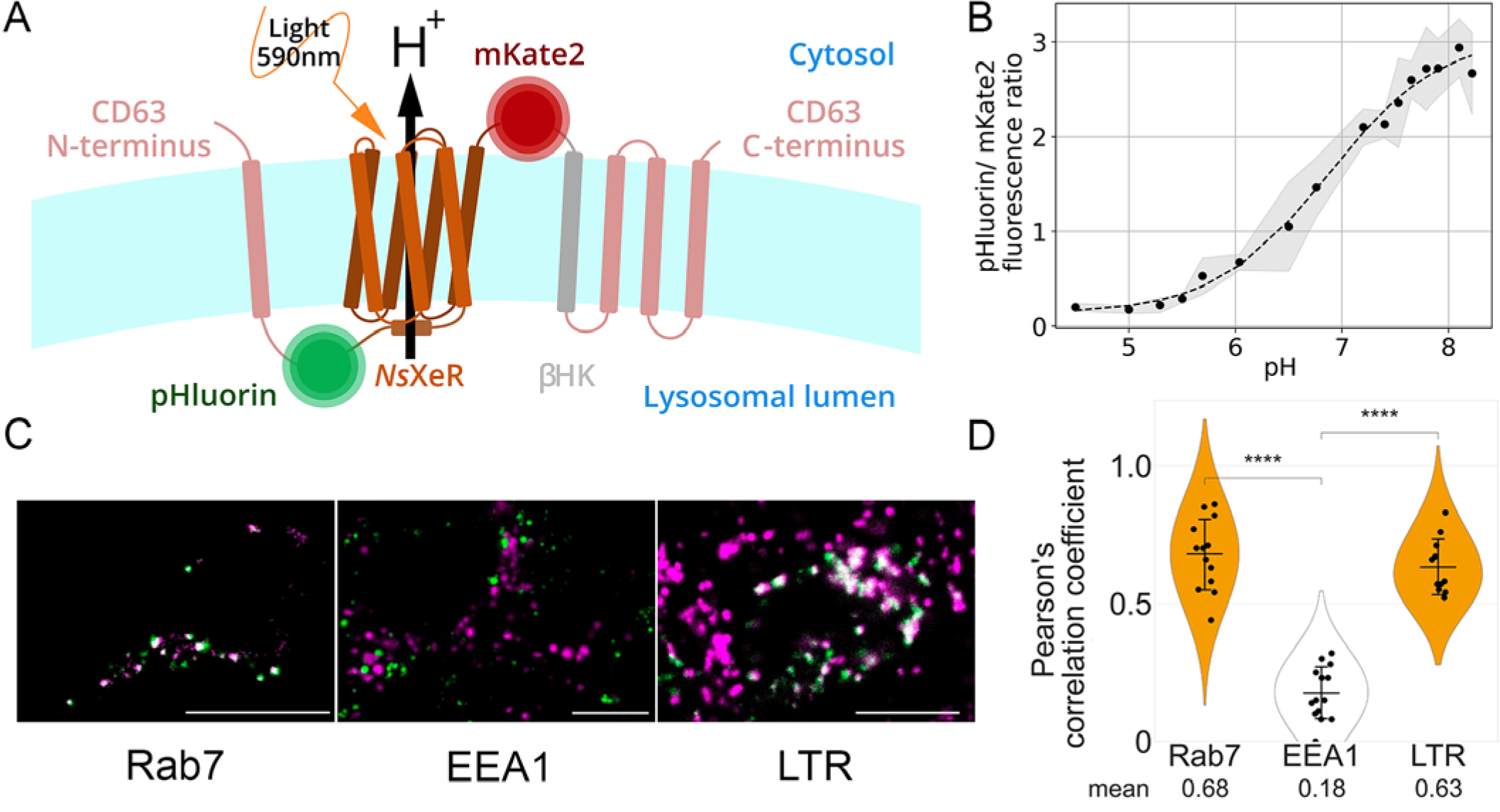
Lyso-*Ns*XeR tool representation. (A) Schematic representation of the lyso-*Ns*XeR (CD63N-se-pHluorin-*Ns*XeR-mKate2-βHK-CD63C) construct consisting of CD63 alpha-helixes flanking *Ns*XeR, the fluorophores (pHluorin and mKate2), and the βHK transmembrane domain. (B) Calibration plot for the convertation of pHluorin/mKate2 fluorescence ratio to lysosome pH. Data represented as mean±SD, Data collected from 5 alive HEK293T cells. (C) Preferential lysosome targeting of lyso-*Ns*XeR construct. Color code: violet – Rab7/EEA1/LTR, green - lyso-*Ns*XeR. Scale bars 10μm. (D) Quantification of overlap between mKate2 (lyso-*Ns*XeR construct marker) and organelles markers. Data represented as mean±SD, N cells >10. Student’s t-test (independent samples, with Bonferroni correction) was used to test for statistical differences in the data with normal distribution, ****: p-value ≤ 10^−4^.

Lyso-*Ns*XeR construct allows to monitor pH inside the lysosomes using pHluorin/mKate2 fluorescence ratio and calibration curve (Figure 2B).

The lyso-*Ns*XeR lysosomal localization verification presented in Figures 2C, D. Fluorescence confocal microscopy in the HEK293T cells transfected with lyso-*Ns*XeR shows that mKate2 (red fluorescence) from the lyso-*Ns*XeR construct is co-localized sufficiently with Rab7, protein marker of late endosomes/lysosomes, and LysoTracker Red (orange fluorescence), staining acidic lysosomes, whereas co-localization with early endosomes marker EEA1 is poor.

The lyso-*Ns*XeR ability to alkalize lysosomes via xenorhodopsin *Ns*XeR proton pumping presented in Figure 3. An increase of pHluorin fluorescence in the lyso-*Ns*XeR transfected but chemically untreated HEK293T cells (Figure 3A, 5 regions of interest, ROIs) is observed under illumination with 590nm light (Figure 3B, green curve). The undesirable effects linked to the lysosomal movement during imaging are not present since mKate2 red fluorescence does not change significantly (Figure 3B, red curve); whereas pHluorin fluorescence is noticeably and synchronously increased under illumination and decreased without it. Therefore, the pHluorin fluorescence changes reflect pH alteration rather than the lysosome movement. As expected, the emerging pH change produced by lyso-*Ns*XeR depends on the light intensity, increasing up to saturation under higher LED power values (Figure 3C).

**Figure 3.**
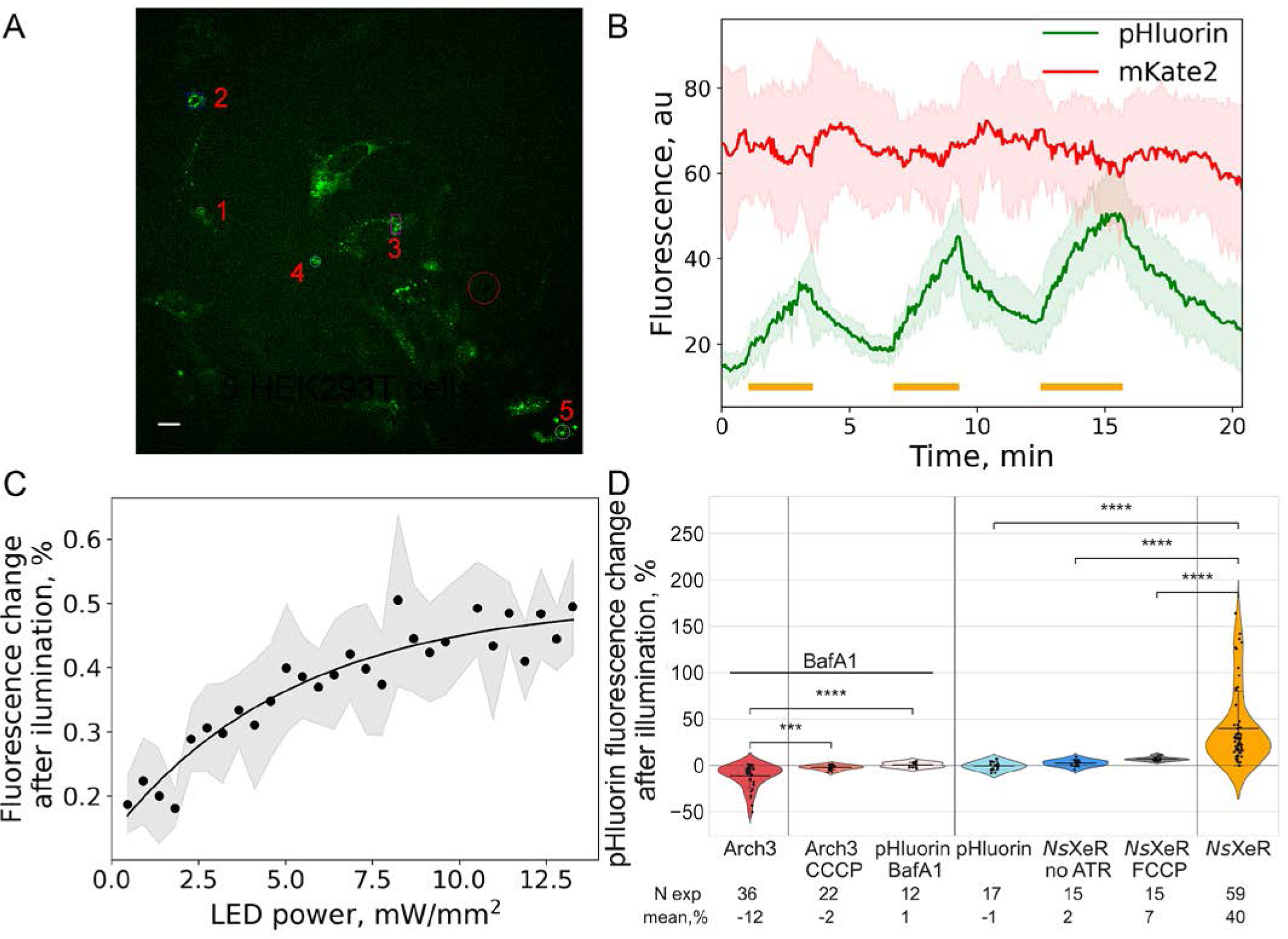
Optogenetic de-acidification of lysosomes in HEK293T cells by Lyso-*Ns*XeR. (A) Confocal image of HEK293T cells, transfected with lyso-*Ns*XeR. 1-5 numbers show cells regions of interest (ROIs) in the image, which were used for fluorescence changes collection. Scale bar 10μm. (B) Demonstration of lysosomal de-acidification (by pHluorin sensor, green curve) under cells illumination by LED 590nm light, activating *Ns*XeR proton pumping. The absence of mKate2 fluorescence change (red curve) proves that lysosomes are not moving during observation. Periods of illumination are highlighted by orange lines. The fluorescence signals were corrected on background to remove backlight from LED. Time curves presented as mean (for the data from 5 ROIs) ± SD. See also Supplementary Movie 2 for ROI 2. (C) The dependence of the lyso-*Ns*XeR light-induced pHluorin fluorescence change on illumination intensity (LED power). The effect plateaued after exponential growth. Curve presented as mean (for the data from 5 cells) ± SD. (D) Control experiments with lyso-Arch3 and lyso-*Ns*XeR constructs in HEK293T cells. Conditions: lyso-Arch3 (with overnight 50nM BafA1), CCCP: 50uM 1h, lyso-*Ns*XeR. FCCP: 50uM 1h. Choose of FCCP as stronger protonophore for lyso-*Ns*XeR was only effective in the case of untreated cells. Mann-Whitney-Wilcoxon test (two-sided, with Bonferroni correction) was used to test for statistical differences in the data without normal distribution, ***: 10^−4^ ≤ p-value ≤ 10^−3^, ****: p-value ≤ 10^−4^. N corresponds to the number of the light shots in all experiments.

Our control experiments (Figure 3D) with the construct lacking rhodopsin (lyso-pHluorin) and lyso-*Ns*XeR with protonophore FCCP or in absence of all-trans retinal (ATR, rhodopsins cofactor) show that the main reason for the observed pHluorin fluorescence change is *Ns*XeR light sensitivity. The protonophore diminishes the lyso-*Ns*XeR induced pHluorin fluorescence increase (pH change) equilibrating proton concentrations inside and outside lysosome (Figure 3D). The same control experiments were done for lyso-Arch3 construct showing the absence of significant effects on pHluorin fluorescence under illumination of the lyso-pHluorin construct (without rhodopsins) with BafA1, under illumination of the lyso-Arch3 construct in the presence of protonophore (Figure 3D).

For higher accuracy, it should be pointed out that the control experiments with the lyso-pHluorin (without rhodopsin) construct in BafA1-treated and untreated cells showed a slight change of the amplitude of pHluorin fluorescence upon the maximum power of 590 nm illumination (Figure 3D). However, this effect is significantly lower than the pH change triggered by lyso-*Ns*XeR and lyso-Arch3 and should not complicate interpretation of the data obtained in the optogenetic experiments.

We measured in absolute units the maximum lysosome pH increase triggered by light activated lyso-*Ns*XeR. The pHluorin fluorescent changes were normalized on the protein expression by measuring the ratio of pHluorin to mKate2 fluorescence. This ratio determined at different lysosomal pH allows calibrating the pH-sensor and calculating ΔpH in absolute units from pHluorin rises after light activation of the rhodopsins. The calibration plot (Figure 2B), which agrees well with the literature data for pHluorin (Sankaranarayanan et al., 2000), shows that lyso-*Ns*XeR increases pH up to 1 unit under illumination (Figure 4A).

**Figure 4.**
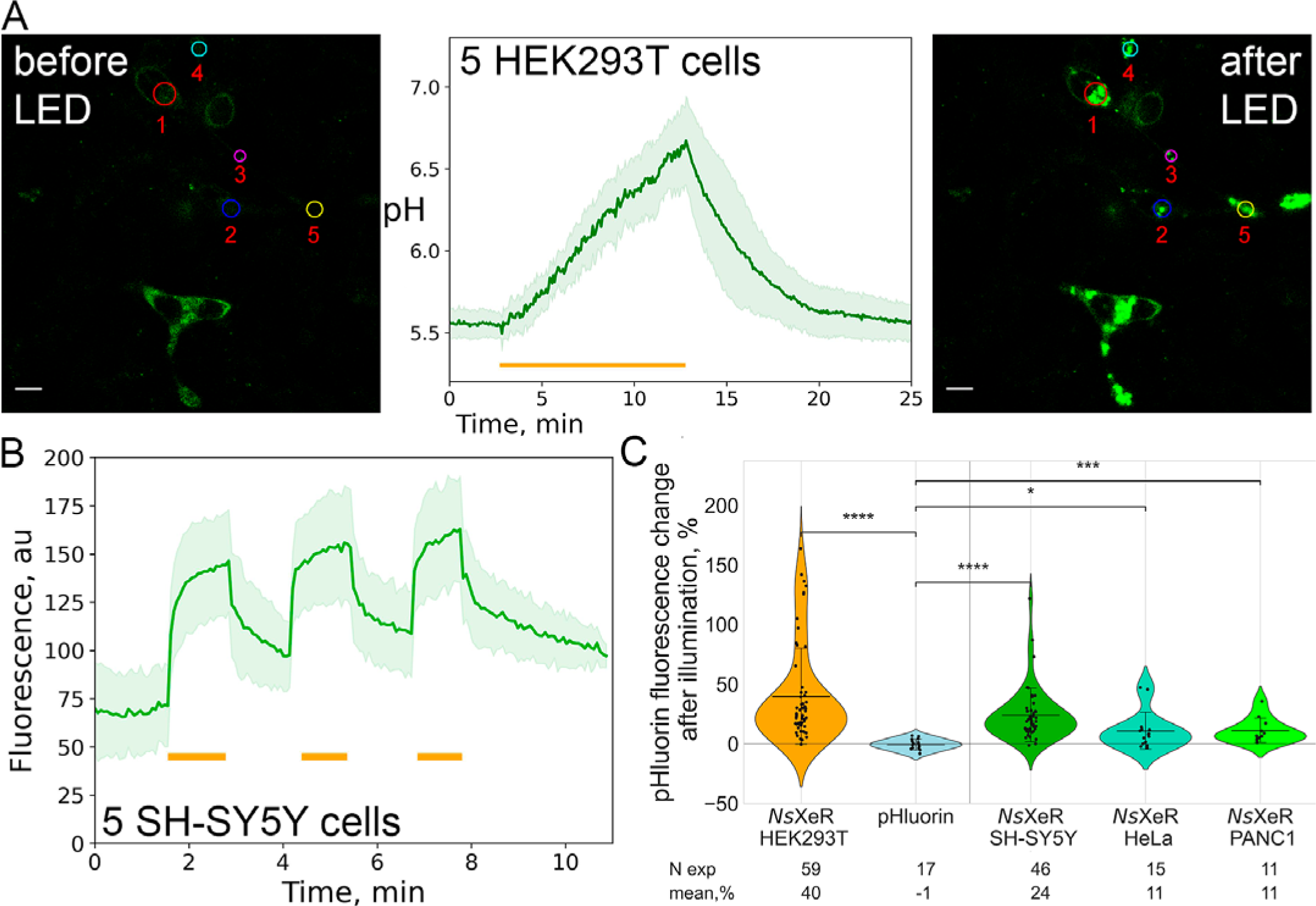
Lyso-*Ns*XeR validation: ΔpH and different cell lines. (A) Demonstration of maximum lysosomal alkalization in HEK293T cells under prolonged illumination, pHluorin fluorescence converted to pH by calibration plot (Figure 2B). The pH change about 1 unit reveals high potency of the approach. See also Supplementary Movie 3. (B) Light-driven lysosomal pH increase by lyso-*Ns*XeR expressed in SH-SY5Y neuroblastoma cell line. Periods of illumination are highlighted by orange lines. The fluorescence signals were corrected on background to remove backlight from LED. Time curves presented as mean (for the data from 5 ROIs) ± standard deviation (SD). (C) Validation of pHluorin changes in SH-SY5Y, HeLa, PANC1 cancer cell lines. Mann-Whitney-Wilcoxon test (two-sided, with Bonferroni correction) was used to test for statistical differences in the data without normal distribution, *: 0.01 ≤ p-value ≤ 0.05, ***: 10^−4^ ≤ p-value ≤ 10^−3^, ****: p-value ≤ 10^−4^. N corresponds to the number of the light shots in all experiments.

Thus, the combined lyso-*Ns*XeR and lyso-Arch3 optogenetic system would allow controlling lysosomal pH in the widest possible pH range within two units. We expect that further optimization of the proteins and their expression in lysosomes may further increase the range. Considering high sensitivity of the cell and, in particular, lysosome functions to pH changes, we also expect that lyso-*Ns*XeR itself and in combination with lyso-Arch3 provides powerful optogenetic approach to the studies of the pH dependent functions and dysfunctions of lysosomes.

Notably, the pH-sensing fluorescent protein pHluorin is located in the vicinity of *Ns*XeR, but collected fluorescence change curves reflect pH of the bulk phase of lysosomal lumen due to fast H^+^ diffusion in water volume (diffusion coefficient is ∼10^4^ µm^2^/s (Agmon, 1995)). Having in mind that lysosome diameter is ∼0.5µm (de Araujo et al., 2020), so its volume is ∼10^−17^ l, under pH 4.5, 5.5, 6.5 proton amount inside the lysosome lumen is ∼400, 40, 4 protons, correspondingly. Thus, proton pumping by *Ns*XeR cause firstly fast local ‘surface’ juxta-membrane effect of alkalization and pHluorin fluorescence increase, following proton diffusion out of lumen and pumping out, but, under used experimental setup, we assume that these steps could not be separated within monitoring with the period of ∼4s.

Amplitude and kinetics of light-induced lysosome de-acidification depends, on the one hand, on the rhodopsin expression level and, on the other hand, on acidification by vATPase, which also has heterogeneity by the cellular state and the localization of lysosomes in the cells (Johnson et al., 2016). Additionally, endogenous proton leaks also occur (Xu and Ren, 2015). Our experiments were performed without inhibition of vATPase so expressed lyso-*Ns*XeRs can pump protons probably more effectively than endogenous lysosomal vATPase allowing increase of lysosomal pH in the untreated cells.

The proposed here approach allows time-resolved monitoring of re-acidification (relaxation of intraluminal acidification) kinetics just after turning off the illumination of the cells. Since *Ns*XeR proton pumping terminates within 30 ms (the length of photocycle) (Shevchenko et al., 2017) after the light is switched off, post-illumination relaxation shows pure endogeneous pH adaptation that could be mainly attributed to vATPase working capacity (Riederer et al., 2023), with some impact of other acidifiers (Fujii et al., 2023). Further experiments should be done for clarification. Altogether, coupled pH modulation and detection is a unique *in situ* approach to lysosome pH equilibrium adjustment investigation in terms of simplicity and pithiness.

To test ability of lyso-*Ns*XeR to increase pH in lysosomes of cancer cells we repeat experiments in SH-SY5Y, HeLa, PANC1 cancer cell lines (Figure 4B, C). Data prove applicability of lyso-*Ns*XeR to impact not only model but real cancer cells, showing the possibility of widespread application of the proposed approach.

Post-illumination re-acidification takes minutes (look Figure 4A); therefore one could analyze lyso-*Ns*XeR assisted lysosomes de-acidification also by flow cytometry to obtain data from many cells. The flow cytometry data confirmed the pronounced effect of lyso-*Ns*XeR on the intralysosomal pH. HEK293T cells transfected by lyso-*Ns*XeR show lysosome fluorescence increase in green (pHluorin) channel without change in red (mKate2) channel after illumination in comparison with non-illuminated cells (Figure 5A). Histograms overlay for the pHluorin/mKate2 ratio (Figure 5B) and comparison of effects in repeats (Figure 5C) show that *Ns*XeR-dependent alkalization under illumination could be similar to the alkalization effect of vATPase inhibition by BafA1, *i.e.,* up to pH ∼6.5 (Yoshimori et al., 1991), which is consistent with microscopy pH change evaluation (Figure 4A).

**Figure 5.**
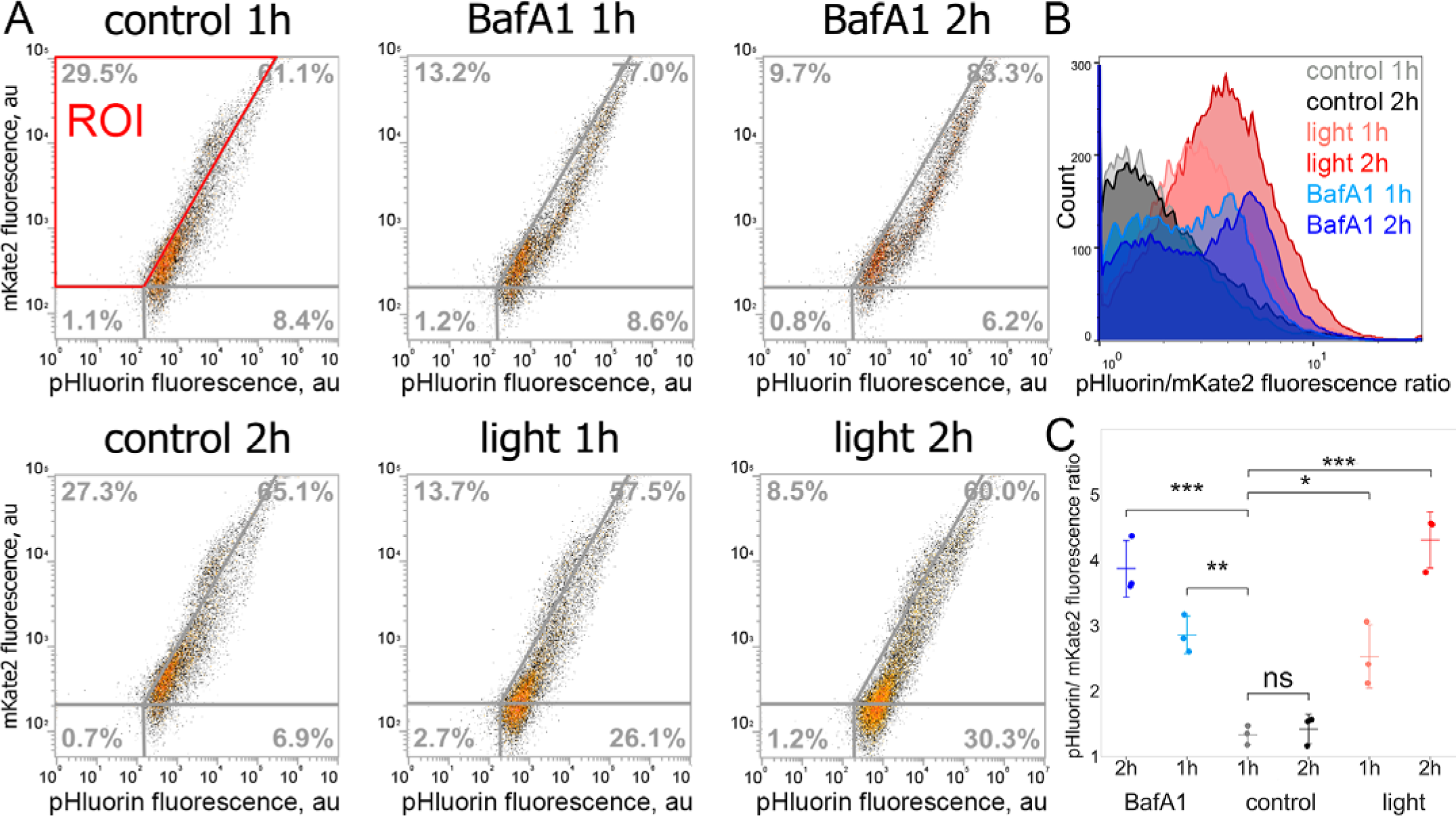
Flow-cytometry confirmation of lyso-*Ns*XeR effectiveness. (A) The representative flow cytometry data comparing BafA1 and lyso-*Ns*XeR effects on lysosomal pH in the lyso-*Ns*XeR-expressing cells, control – lyso-*Ns*XeR expressing cells without treatments. Cells were treated with light or BafA1 and analyzed for a shift in pHluorin/mKate2 ratio, revealing average lysosomes pH per cell, at different time points of treatments. Scatter plots of pHluorin/mKate2 ratio are shown with Demarcation line made such as in the control sample almost all transfected cells with high mKate2 signal were to the left of the straight line (in the ROI – region of interest). Number of cells in ROI (representing normal acidification of lysosomes) written under scatter plots and show population shift to de-acidified lysosomes. Data collected from more than 10000 alive HEK293T cells. (B) Histograms overlay for the pHluorin/mKate2 ratio showing increase of lysosome pH after BafA1 and lyso-*Ns*XeR action. (C) pHluorin/mKate2 ratio, revealing average lysosomes pH per cell, in transfected cells with high mKate2 signal. Data presented as mean ± SD from three independent experiments. Student’s t-test (independent samples, with Bonferroni correction), ns (non-significant): 0.05 < p ≤ 1, *: 0.01 < p ≤ 0.05, **: 0.001 < p ≤ 0.01, ***: 0.0001 < p ≤ 0.001.

### Justification of the optogenetic control of bulk pH and functional properties of lysosomes: lysosome proteases inactivation by means of lyso-*Ns*XeR

Our optogenetic approach is indispensable for fundamental studies of lysosome functioning and those cell functions and dysfunctions that are coupled with lysosomes. We used Magic Red CTSB cleavage assay (Figure 6A) to show Cathepsin B inactivation by lyso-*Ns*XeR lysosome de-acidification (Figure 6B). The Magic Red CTSB is a reagent exhibiting red fluoresce upon cleavage by Cathepsin B enzyme. Lysosomal cysteine proteases, a subgroup of the Cathepsin family, are critical for normal cellular functions such as general protein turnover, antigen processing, and bone remodeling. Cathepsin B functions in lysosomes for protein degradation and maintenance of cellular homeostasis (Brix et al., 2008). Cathepsin B normally has highest activity at acidic pH in lysosomes. It also cleaves small molecule substrates like Magic Red CTSB at the carboxyl side of Arg-Arg bonds (Cui et al., 2019). In our experiments, lyso-*Ns*XeR transfected and non-transfected HEK293T cells were loaded by Magic Red CTSB (MR-CTSB) substrate during illumination (Figure 6A). Light application increases pH in lysosomes of transfected cells, detected by pHluorin fluorescence rise with no significant change in non-transfected ones (Figure 6C, D, green and blue curves go up synchronously after MR-CTSB addition dueo to focus change). In turn, in red fluorescence channel, where cleaved MR-CTSB emits, non-transfected cells show around two-times higher signal plateau than lyso-*Ns*XeR transfected ones (Figure 6E, F). Proved by repeats (Figure 6B), lyso-*Ns*XeR optogenetics enables unprecedented time resolution control of the activity of lysosomal enzymes.

**Figure 6.**
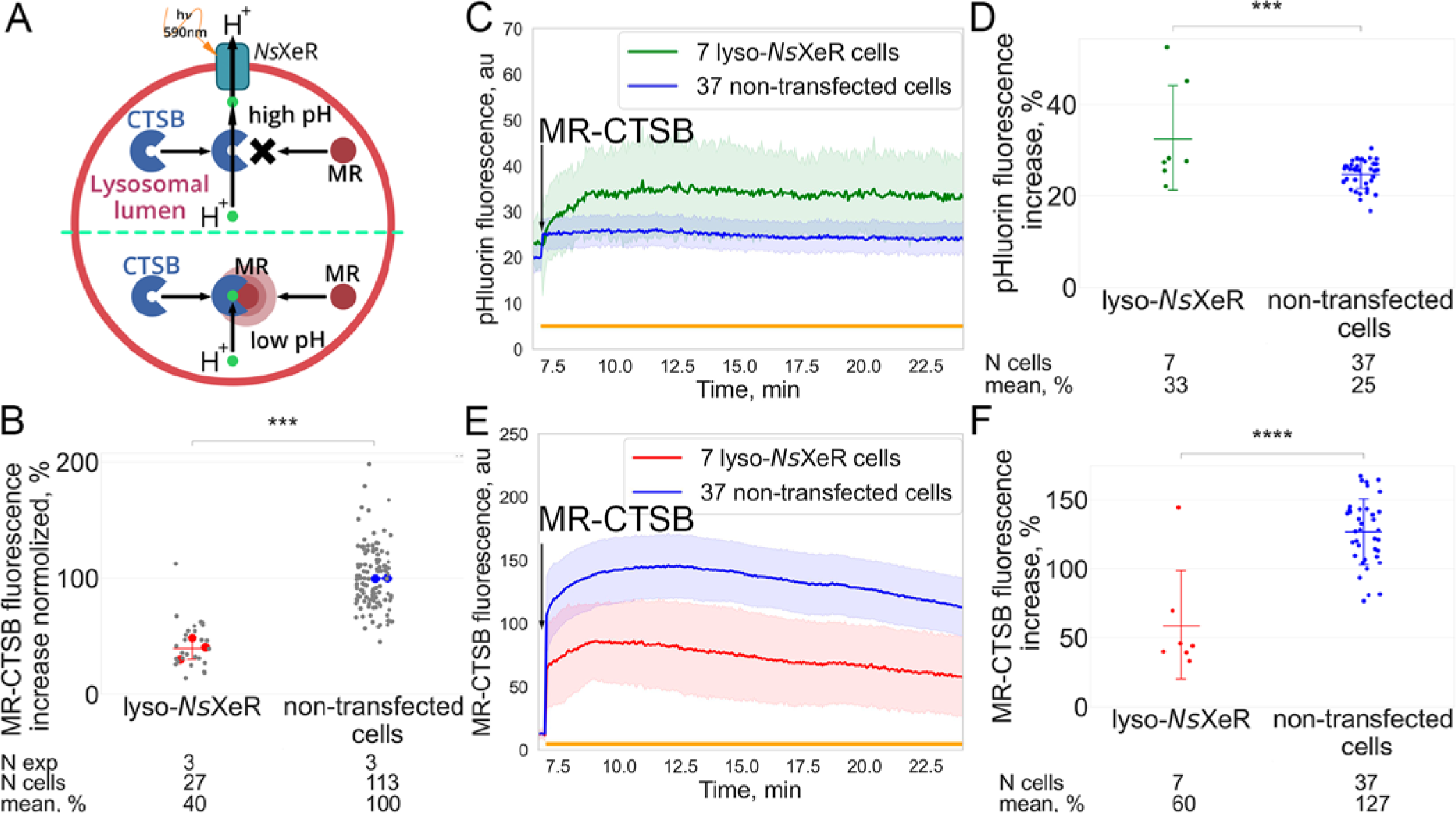
Activation of lyso-*Ns*XeR increases pH of lysosomal lumen and inhibits proteases in HEK293T cells. (A) Magic Red CTSB cleavage assay: Cathepsin B (CTSB) cleaves Magic Red-CTSB substrate (MR here) in the lysosomes with low pH and red fluorescence arise. Lyso-*Ns*XeR de-acidifies lysosomes and therefore inhibits CTSB activity, so there is no so high increase of red fluorescence. (B) Lyso-*Ns*XeR-mediated optogenetically induced pH increase inhibits lysosomal proteases that is revealed by difference in red fluorescence increase in 3 repeats between 27 transfected and 113 non-transfected cells. (C) Light activation of lyso-*Ns*XeR increases pHluorin fluorescence, *ie* lysosome pH. (D) Quantification of pHluorin fluorescence increase after illumination in transfected and non-transfected cells. (E) Light activation of lyso-*Ns*XeR makes increase of red MR-CTSB fluorescence lower, *ie* inhibits CTSBs. (F) Quantification of MR-CTSB fluorescence increase after illumination in transfected and non-transfected cells Data presented as mean ± SD. Student’s t-test (independent samples, with Bonferroni correction), ***: 0.0001 < p ≤ 0.001, ****: p ≤ 0.0001 Time curves presented as mean (for the data from specified number of cells) ± standard deviation (SD).

## Discussion

In the present work we introduced a new approach and demonstrated that important cell functions can be controlled simply by light through modulation of lysosomal pH with unprecedented spatial and time resolution. It is shown that it is a highly efficient approach to the studies of lysosome associated functions and dysfunctions of cells. Expression of inward light driven proton pump (*Ns*XeR) in lysosomes is highly selective, and this provides outstanding spatial resolution of the approach. Complementary use of lyso-*Ns*XeR and lyso-Arch3 can vary lysosome proton concentration by about two orders in both directions (alkalization (Figure 1C) and acidification (Figure 4A)).

Lysosome alkalization by lyso-*Ns*XeR is unique by its selectivity of action in contrast with other vATPases inhibiting drugs. For example, BafA1 blocks vATPases in lysosome and cellular membranes (Forgac, 2007) and furthermore inhibiting Ca^2+^ pump SERCA (P-ATPase) in endoplasmatic reticulum (Mauvezin et al., 2015). Concanamycins also binds with v- and P-ATPases (Dröse and Altendorf, 1997).

Moreover, simultaneous compensation or exacerbation of the possible cytosolic acidification, produced by lyso-*Ns*XeR, could be done by usage of rhodopsin proton pumps, targeted to plasma membrane. Thus, pure consequences of lysosomal alkalization could be investigated by lyso-*Ns*XeR system.

To make clear whether lysosome optogenetics is indeed able to control the key lysosomal functions, we demonstrated the lyso-*Ns*XeR optogenetic regulation of Cathepsin B activity, which provides protein degradation and supports cellular proteostasis.

The fact that proton concentration can be controlled via lyso-rhodopsin proton pump optogenetics in a range of at least two orders of magnitude is encouraging for the extension of the proposed approach to other rhodopsin ion pumps (chloride, sodium, etc.). Indeed, rhodopsin ion pumps have a comparable length of photocycle and therefore similar ion pumping efficiency. Passive permeability of biological membranes to protons is usually considerably higher than to other ions. It opens new perspectives in lysosome and cell studies. Lysosomes are significantly enriched with chloride, and a decrease of chloride concentration correlates directly with a loss in the degradative function of the lysosome (Chakraborty et al., 2017; Graves et al., 2008). A decrease of lysosomal chloride influences Ca^2+^ release from the lysosome and impedes the activity of specific lysosomal enzymes, indicating a broad role for chloride in lysosomal function (Chakraborty et al., 2017).

Considerable variation of ion concentration is possible due to a high ratio of the membrane surface to the volume of lysosomes determined by a small size of the organelle. The size of other cell organelles is also small in comparison to the size of the cells. It means that organelle optogenetics based on ion transporters can control many cell processes through light modulation of different ions concentrations. In future, lysosome optogenetics could be translated to lysosome-related organelles (Saftig and Klumperman, 2009) and other organelles for the studies of cellular functions and dysfunctions dependent on ions concentration (Thattai, 2017; Weisz, 2003). It encourages extension of the organelle optogenetic toolkit with other ion pumps and channels.

Optogenetic control of the lysosome function through pH regulation over a wide range allows studying a wide range of fundamental lysosome depending cell functions, including lysosomal-mitochondrial axis, lysosome-to-nucleus signaling, ER remodeling, metabolic adaptation and rewiring, and virus egress. The introduced lysosome optogenetics may find a range of applications in the studies of the diseases associated with the dysfunction of lysosomes, like lysosomal storage diseases (Parenti et al., 2021; Platt et al., 2018). The analysis of possible applications of lysosome pH control was carried out in the Supplementary Note 1. One might think that lysosomal alkalization has only pathogenic impact. It is tempting to apply lyso-*Ns*XeR tool to kill diseased cell, for example overactivated autoimmune cells in autoimmune disorders (Cao et al., 2021), influencing specific tumor features (Bonam et al., 2019), relying on lysosome over-acidification (autophagy addiction (Levy et al., 2017), invasion (Hämälistö and Jäättelä, 2016), cytosolic alkalization (Webb et al., 2011)).

Alternative resembling approaches for light-controlled cellular toxicity like photodynamic cancer therapy using chemical photosensitizers is known (Li et al., 2020). Feasible advantage of optogenetical impact on cancer is the possibility of using selective expression (e.g. by hypoxia-specific promoters (Brown and Wilson, 2004), or exploiting of transcriptional addiction (Vervoort et al., 2022)).

Lysosome acidification is a mainstream among anti-aging interventions (Aman et al., 2021). However, there is possible advantage of transient lysosome alkalization over acidification. Senescent cells have alkali lysosomes due to their membrane permeabilization (Johmura et al., 2021), so acidification is senseless at this stage. On the contrary, transient alkalization on pre-senescent cells could initiate lysosome biogenesis by TFEB activation (Lie and Nixon, 2019), delaying process of autophagy decay.

## Methods

### Reagents

Bafilomycin A1 (BafA1) (B1793), Carbonyl cyanide 3-chlorophenylhydrazone (CCCP) (C2759), Carbonyl cyanide 4-(trifluoromethoxy)phenylhydrazone (FCCP) (C2920) were from Sigma (Merck). Cathepsin B Assay Kit (Magic Red CTSB) (ab270772) was from Abcam. LysoTracker Red DND-99 (L7528), pHrodo Red AM (P35372), Phusion™ High-Fidelity DNA Polymerase (F530), FastAP Thermosensitive Alkaline Phosphatase (E0651), FastDigest NotI (FD0594), FastDigest KpnI (FD0524) were from ThermoFisher Scientific. Polynucleotidekinase T4 (E311), DNA ligase T4 (E319) were from SibEnzyme. Plasmid MiniPrep (BC021S) and CleanUp Standart (BC022) were from Evrogen, Russia.

### Genetic constructs creation

pCMvlyso-pHoenix and pCMvlyso-pHluorin were a gift from Christian Rosenmund (Addgene plasmid # 70112, 70113), pTagBFP-Rab7, TagRFP-EEA1 were used for colocalization studies. The lyso-*Ns*XeR constructs were made by followed procedure. In the pCMvlyso-pHoenix, plasmid BsiWI was replaced with the NotI restriction site by PCR. Afterwards, the *Ns*XeR gene (Shevchenko et al., 2017) was cloned by restriction-ligation with the KpnI & NotI sites. Lyso-*Ns*XeR construct with TagBFP pH-insensitive fluorescent protein sequence was obtained by replacement of mKate2 to TagBFP. Cloning was approved by DNA sequencing.

### Cell cultivation, transfections and treatments

Human cells HEK293T (ECACC 12022001) were cultivated in DMEM (Gibco, Cat.No: 41966) in CO2-incubators (Binder C150 E2) at 37°C, 5% CO2 and high humidity. For microscopy experiments, cells were seeded on the ibitreated 35 mm dishes (ibidi, Cat.No: 81156). Transient transfections were done on these dishes using 2.5ug plasmid and 6ul Lipofectamine 3000 (Thermo Fisher L3000008) two days before imaging. All-trans retinal was added to the final 10uM concentration to the cells growth medium simultaneously with transfection. For the lyso-Arch3 experiments, the HEK293T cells were treated with 50 nM BafA1 overnight for full lysosome alkalization. For the lyso-*Ns*XeR experiments, untreated HEK293T cells were used.

### Optogenetic experiments

Live-cell imaging was done, as previously described (Bogorodskiy et al., 2021). We used an inverted confocal LSM780 microscope (Carl Zeiss, Germany), 63x (NA = 1.4, oil immersive) objective. Optical fiber (400 um diameter) guided through a microinjection needle holder was placed just above the cells of interest by means of a microinjection micromanipulator InjectMan NI2 (Eppendorf). LED590 (ThorLabs M590F2, 590nm emission maximum) was used with a power of 16 mW/mm2 (ThorLabs Thermal Power Sensor Head S302C measured) in all experiments, excluding the ones with LED power variation (ThorLabs 4-Channel LED Driver, DC4104). During the experiments, the cells were placed in a Tokai Hit CO2-incubator (model INUBG2H-ELY), HEPES (pH 7.4) for the final concentration 25 mM was added. For time series with light illumination, two approaches were used: a channel mode and a *λ*-mode.

Channel mode. First, for the control experiments, excitation at 488, 561 nm was used respectively for the simultaneous detection of mKate2 and pHluorin fluorescence for lysosome movement accounting. The emission was measured in CLSM Channel-mode using a 2-channel detector (Carl Zeiss, Germany) set to a 488–523 nm (Channel 1) and 635–712 nm ranges (Channel 2) to avoid detection of maximum intensity of LED590. Slight LED590 emission was passed to Channel 1 and Channel 2 and was used to indicate time when LED590 worked. Time series experiments were done with time for one image obtaining 3.88s with no interval between the images, so pHluorin intensities were collected with 3.88s intervals. Time curves were obtained using ZEN software by Zeiss.

LED590 emission, which added intensity to Channel 1 or 2, was obtained from background ROI (near with lysosomes ROI) and then was subtracted from the time curves of lysosomes ROI to produce pure pHluorin and mKate2 fluorescence intensity time curves, correspondingly.

*λ*-mode. pHluorin fluorescence intensity only was excited by a 488 nm laser. The emission was measured in CLSM *λ*-mode using a 34-channel QUASAR detector (Carl Zeiss, Germany) set to a 488–535 nm range to avoid LED590 maximum intensity detection. Time curves were obtained using ZEN software by Zeiss after performing spectral Linear unmixing (in ZEN software by Zeiss) processing option (GFP spectrum assignment for spectrum of each pixel was done).

For Figure 2A, pHluorin fluorescence relative change were automatically obtained from the time series for the each illumination. Each pHluorin fluorescence relative change value is calculated for the full field of view (image) with many cells and lysosomes in each. Video files obtained were converted to png images by ZEN software by Zeiss. Python code (Supplement Notebook) did data array for all images of each experiment. The masks for cells, lysosomes and extracellular background for the each image were found using min max filter (Verbeek et al., 1988). Average intensity in pixels of the lysosome mask was found for each image and average intensity of the extracellular background (reflecting LED illumination) was subtracted. Curves of fluorescence changes in time for lysosomes were generated. Optogenetic effect was calculated as (Ff-Fi)/ (Ff+Fi)*100%, where Ff and Fi correspond to pHluorin fluorescence intensity right after finishing the illumination and before illumination, respectively. Tukey’s post hoc tests were done to calculate a P value.

For Figure 3D pHluorin fluorescence change calculated as |(Ff-Fi)/Fi|*100%.

LED590 emission, which added intensity to the GFP spectrum, was obtained from background ROI and then was subtracted manually from the time curves of lysosomes ROI to produce pure pHluorin fluorescence intensity time curves. Finally, the images were aligned and positioned using Adobe Photoshop (Adobe Systems, USA).

### pH measurements in lysosomes

HEK293T cells were incubated with different pH buffer solutions (based on HBSS, Gibco, cat No 14170) in the presence of Digitonin (10 µM) and Bafilomycin A1 (100 nM). For calibration by the GFP/RFP ratio (pH from 5.5 to 8.5) images from five cells in three different dishes were imaged in *λ*-mode (488–700 nm range), and Unmixing option of ZEN software by Zeiss was used for spectrum generation. For quantification of the lysosome acidity, the GFP/RFP fluorescence ratio was measured after excitation by a 488 nm laser - 0.4%, 561nm - 0.1%, because of 6-fold lower power of 488nm laser but 1,5-fold higher quantum yield of pHluorin (GFP) compared to mKate2 (RFP). Then the ratios of fluorescence intensities of pHluorin at 510nm (GFP value) and mKate2 at 620nm (RFP value) were calculated from the spectrum generated for the Region of Interest by Unmixing option in ZEN software by Zeiss.

### Colocalization study

HEK293T cells were transfected by lyso-*Ns*XeR construct and with markers of organelles of interest: Rab7-BFP: (late endosomes/lysosomes); TagRFP-EEA1: Early Endosome Antigen 1, marker for early endosomes; LTR: LysoTracker Red DND-99 (acidic orgenelles).

Pearson coefficient for co-existence of fluorescent protein in marker protein (BFP for Rab7, TagRFP for EEA1) and pH-insensitive fluorescent protein lyso-*Ns*XeR construct (mKate2 or BFP correspondingly) was obtained in Coloc option in ZEN software by Zeiss for *λ*-mode images after Unmixing.

### Cytometry

Cell analysis on stable alkalization was done on an Attune NxT flow-cytometer (Applied Biosystems, Thermo Fisher) using BL1 (488nm laser excitation, 512/25nm detection), YL2 (561nm laser excitation, 620/15nm detection) channels for pHluorin and mKate2 fluorescence detection, correspondingly. Light illumination of the detached cells in a 1,5ml tube was done using Olympus KL1500-LCD (250W) with a 605/70nm optical filter. A two-parameter display of FSC vs. SSC was set up excluding subcellular debris. Single cells were selected using a two-parameter display of FSC-H vs. FSC-A. Such cells were used for further fluorescence analysis.

### Live imaging of Magic Red CTSB staining

Magic Red (Arg-Arg-Cresyl Violet-Arg-Arg) (Abcam), fluoregenic substrate, emitting red fluorescence after Cathepsin B cleavage, was added to the HEK293T cells during the Time Series experiment directly to 2 ml medium. Vial with dry substrate for 50 reactions was dissolved in 50ul DMSO. This stock solution was diluted 1:10 with UltraPure Water (10977-035, Invitrogen) and added to the cells in the amount of 5-10ul depending on the cells quantity on the imaging dish. The Rate of Magic Red CTSB cleavage, revealed by the appearance of red fluorescence, is slower in alkali lysosomes than in acidic ones, which should be another proof of possibility of lyso-*Ns*XeR to alkalize lysosomes.

### Statistics

No statistical methods were used to predetermine sample size. Microscopy results are representative images, Pearson correlation coefficients were obtained for the assessment of colocalization from at least 10 cells. Cytometry experiments were conducted at least in n=3, in each repetition more than 10000 cells were tested. The Python libraries scipy, statannot were used for statistical analysis. All data are presented as mean ± SD (lines), Student’s t-test with Bonferroni correction was used to assess statistical significance, p-value annotation was done as: ns: 0.05 < p <= 1, *: 0.01 < p <= 0.05, **: 0.001 < p <= 0.01, ***: 0.0001 < p <= 0.001, ****: p <= 0.0001.

## Abbreviations

Arch3: archaerhodopsin from *Halorubrum sodomense*
*Ns*XeR: xenorhodopsin from the nanohalosarchaeon Nanosalina
vATPase: Vacuolar-type H^+^-ATPase, enzyme responsible for maintaining lysosomal pH by ATP-dependent proton pumping
TFEB: transcription factor EB
ER: endoplasmatic reticulum
HEK293T: Human Embryonic Kidney 293 with expression of mutant version of the SV40 large T antigen cell line for immortalization through p53 inhibition
BafA1: Bafilomycin A1, vATPase inhibitor
pCMV: expression vector with Cytomegalovirus promoter
CD63: Cluster of Differentiation 63
βHK: β-subunit of H+,K+-ATPase
Rab7: Rab7 protein, marker for late endosomes/lysosomes
EEA1: Early Endosome Antigen 1, marker for early endosomes
LTR: LysoTracker Red DND-99, marker of acidic orgenelles
ATR: all-trans retinal
lyso-: lysosomal
SD: standard deviation
ROI: region of interest
LED: light emitting diode
GFP: green fluorescent protein
se-pHluorin: super-ecliptic pH-sensitive GFP mutant pHluorin
RFP: red fluorescent protein
CLSM: confocal light scanning microscope
CTSB: cathepsin B
FCCP: Carbonyl cyanide 4-(trifluoromethoxy)phenylhydrazone
CCCP: Carbonyl cyanide 3-chlorophenylhydrazone
ATP: adenosine triphosphate
SH-SY5Y: human neuroblastoma cell line
PANC-1: epithelioid carcinoma attached cell line that is currently used as an in vitro model to study pancreatic ductal adenocarcinoma carcinogenesis and tumour therapies
HeLa: cervical carcinoma cell line
DMEM: Dulbecco’s modified Eagle’s medium.

## Acknowledgements

We thank E. Shestoperova, V. Schukina, I. Panov, V. Borisov, A. Melnikov for their help with the experiments, Norbert Dencher and Elizaveta Podolyak for their fruitful discussions. The optogenetic setup construction fluorescence was done with support of the Ministry of Science and Higher Education of the Russian Federation (Agreement # 075-03-2023-106, Project FSMG-2020-0003). Lyso-*Ns*XeR optogenetic experiments, including those under conditions of different cell treatments, were supported by Russian Science Foundation (RSF) Project 21-64-00018. The lysosomal enzymes activity tests, the statistical analysis of the experimental data were done with support of the Ministry of Science and Higher Education of the Russian Federation (Agreement # 075-01593-23-04, Project 720000F.99.1.BN62AA40000).

## Author contributions

NI, SB and DB contributed equally, and each has the right to list himself first in bibliographic documents. FT, SB, AV, AM, OM did work on the genetic constructs creation; NI, SB, VA did cell cultivation, transfections, and treatments; VG, NI, SB, SVN planned the experiments design; NI supervised most of the experiments; NI, SB, DB, VA performed the optogenetic experiments, including the study of different responses under various treatments, NI, SB studied the optogenetic impact on the effectiveness of lysosomal enzymes; SB, NI, AB, VB revealed artifacts in the microscopy data; AA, SFN did automatization of image processing; VG, AA, EB supervised the optogenetic experiments; SVN, VG, NI, EB, AR made manuscript conceptualization; DB, NI did the cytometry experiments; VG initiated, provided general supervision of the project and together with NI, SVN and the contribution from all other co-authors who prepared the manuscript.

## Additional information

**Supplementary Information** accompanies this paper at link

