## supplemental information for "Optogenetic control of lysosome function"

#### This PDF file includes:

Supplementary Note 1

References

### Supplementary Note 1. Possible molecular targets of lysosomal acidification and alkalization.

Acidification of lysosomes is convenient intervention for disease treatment and aging delay [11–14]. Another value of lysosomal optogenetics lies in the fact that it is previously lacking technique allowing reversibility and high spatio-temporal resolution for study mechanisms of homeostasis loss due to lysosome alkalization/acidification.

Optogenetic control of the lysosome function through the regulation of pH over a wide range allows studying a wide range of fundamental lysosome depending cell functions, including lysosomal-mitochondrial axis [15], lysosome-to-nucleus signaling [16], ER remodeling [17,18], metabolic adaptation [19] and rewiring [20,21], virus egress [22].

Lysosome alkalization induces cellular responses in different key directions - lysosomal-mitochondrial axis, lysosome-to-nucleus signaling, ER remodeling, metabolic adaptation and rewiring. Attempts of compensation will be fast response, prolonged alkalization will cause successive steps to pathogenesis [12] or cancer inhibition [23]. By using lyso-*Ns*XeR one could uncover benefits and key problems accompanying lysosome alkalization. It should be noted that in comparison to chemical and genetic interventions, fast responses on optogenetic stimulation could be easily revealed (outstanding example is [24]). In contrast, stable prolong impacts will be easier achieved by drugs treatments or genes KD/KO. We describe here possible applications of oprogenetic lysosomal alkalization.

**Calcium signaling, lysosome positioning**

It was recently shown, that pH-dependent lysosomal calcium transporter TRPML1 under vATPase inhibition or lysosome alkalization will induce calcium efflux [25]. In turn, for the some extent lysosome calcium efflux could activate TFEB (if mTORC1 is inactive [16]) and increase lysosome-endosome and lysosome-autophagophore fusion [5]. Lysosome calcium efflux also cause movement **toward to nucleus** [17], their clustering [26] and also supports autophagy.

Thus, transient and reversible optogenetic lysosome alkalization could increase **autophagy** and could be the **fast model for the short acute starvation,** characterized by the same properties (perinuclear lysosome movement, cytosolic alkalization, TRPML1 activation) [19,27]. It also could be directly shown whether TRPML1 activation depends only on luminal pH rather than vATPase activity.

Additionally, mitochondria could receive calcium through lysosome-mitochondria contacts [28], so **mitochondria** also could be influenced by possible lysosomal $Ca^{2+}$ efflux after optogenetic alkalization, resulting in enhanced ATP production efficiency [29]. Importantly, loss of TRPML1 function results in the lysosomal storage disorder mucolipidosis type IV (MLIV) with no lysosomal-mitochondria calcium exchange and several mitochondrial defects [30], so it is tempting to improve lysosome optogenetics tools with calcium transporter.

As lysosome positioning controls **endoplasmatic reticulum** remodeling [18], we suggest that lysosome optogenetic alkalization could have multi-organelle impact.

**Nutrient sensing and metabolic adaptation/circadian rhythms**

Under calorie restriction condition (HBSS) cells have most acidic lysosomes near nucleus [31]. This fact reflects that autophagy is the main source of nutrients for the cells. mTOR activity decreases [27] until the moment of sufficiency of nutrients from autophagic degradation, leading to mTOR activation and autophagic lysosome reformation [32]. mTORC1 is active on peripheral lysosomes in nutrients sufficient conditions [27] and inactive in hypoxic condition [20]. In our study, in full medium (DMEM-FBS, nutrients excess) we see the most acidic lysosomes on the cell periphery, corresponding to the priority of endocytosis and extracellular nutrients source.

**mTORC1** activity is coupled to lysosome amino acids (AA) efflux, partially dependent on vATPase activity (connected to lysosome pH, pH-sensitive AA-transporters) [33], partially on pH-independent SLC38A9 AA-transporter (for review [34,35]). vATPase-Ragulator-Rag complex ("nutrisome") is necessary for the mTORC1 activation, so ligands selectively inhibiting such complex formation by modifying vATPase binding site with no impact on lysosome acidification could be considered as synergistic approach for autophagy activation [36]. Furthermore, lysosome pH controls endocytosis [37], thus acidification is somehow needed for nutrients uptake and sensing in long time periods. Lysosome alkalization produces $Ca^{2+}$ efflux from the lysosome and endoplasmatic reticulum, causing cytosolic $Ca^{2+}$ concentration increase, acting as stress factor that inhibits mTORC1 [38].

**vATPase** is important protein for mTORC1 [3940], AMPK activities [41], onset of senescence [42]. vATPase activity is controlled by reversible dissociation of V0 and V1 subunits. Extent of vATPase association is controlled by concentrations of nutrients such as glucose [43] and amino acids [44]. The first fundamental question needs clarification. There is no clear picture of **vATPase activity change** during metabolic shift from anabolism to catabolism. Just for example, calorie restriction is considering as vATPase activating [4521] or inhibiting [4041] intervention. Only optogenetics with its temporal resolution could reveal each step of metabolic adaptation connected to vATPase.

It is currently commonly accepted that lysosome acidification appears to be dispensable for mTORC1 signaling [3446], only right conformation of vATPase in needed for vATPase-Ragulator complex formation. However, there are arguments that either alkalization or acidification could switch off mTORC1. Chloroquine treatments leading to high lysosome alkalization, without apparent effect on conformation of vATPase, causes mTORC1 inactivation [16]. Furthermore, acidic medium through lysosome movement to cell periphery (probably connected to their acidification and TRPML1 inactivation), cause mTORC1 inactivation [20]. With optogenetically guided lysosome acidifiation and alkalization, uncoupled to vATPase, clarity on the second question could be achieved **- whether lysosomal pH influences mTORC1.**

Inactivation of mTORC1 and consequent TFEB [16] and PGC1a [47] activation could lead to autophagy and lysosome and mitochondria biogenesis. Periodical optogenetical lysosome alkalization could serve as restoration of mTORC1 circadian rhytms, attenuating in amplitude under stress-[20] and aged- [48] conditions.

Third question is could mTORC1 manage vATPase activity? It is known, that mTORC1 regulates vATPase expression [49], as regulator of translation, but whether mTORC1 inhibition could activate vATPase like calorie restriction does (as shown here, Fig. 5d). It is not clear due to position of mTORC1 downstream of vATPase-Ragulator-Rag complex [40].

**ATP depletion,** caused by prolonged lyso-*Ns*XeR activation and enforced vATPase ATP consumption should lead to energetic stress. In case of highly ATP-consuming senescent cells (chemotherapy induced cytostatic cancer cells) it could lead to death and final clearance of unhealthy cells [50].

Thus, lysosome alkalization by lyso-*Ns*XeR optogenetics could reveal different aspects of **nutrient sensing.** While vATPase remains active, lyso-*Ns*XeR-mediated lysosome alkalization, uncoupled from vATPase activity, should inhibit pH-sensitive AA-transporters, so their impact on mTORC1 activity could be shown. Also, dependence of canonical/noncanonical AMPK activity [51], dependent on vATPase [40], could be measured in starvation conditions, while vATPase is inhibited/activated (at the concrete stage of lysosomal adaptation to nutrient starvation), but lysosome is acidic (from lyso-Arch3) or alkaline (from lyso-*Ns*XeR). Such transient and reversible manipulation on lysosomal pH could help discover novel properties of metabolic adaptation [52].

**Cancer cells and stress-response: metabolic rewiring**

Lysosome pH increase prevents iron endocytosis and causes its cellular depletion with subsequent HIF1a stabilization even in normoxia condition [53]. Iron depletion causes mitochondria imbalance and ROS accumulation [15], worsening by inability of mitophagy in alkaline lysosomes (lysosomal-mitochondrial axis). ROS, among other effects, will activate TRPML1 [54], so calcium stress response will be activated in second time (firstly, directly after lysosome alkalization by induction TRPML1 multimerization).

Prolonged lysosome alkalization, iron depletion and HIF1a activation will cause metabolic rewiring like in hypoxia stress or in cancer cells. Glycolysis will acidify cytosol to pH 6.5-7 which suspend circadian clock [20], but cancer cells maintain alkaline cytosol by STAT3-mediated vATPase activation, leading to highly acidic lysosomes [21]. Optogenetic lysosomes alkalization in cancer cells could be the intervention against its invasiveness [5556].

**In future**, lysosome optogenetics could be translated to lysosome-related organelles [57] with possible benefits. Since acidification is required for the function of different cellular organelles[58,59] the lyso-XeR based optogenetics introduced here might have applications beyond lysosomes, for the studies of cellular functions and dysfunctions dependent on the pH state of other organelles due their similar size. It encourages extension of organelle optogenetic toolkit with other ion pumps and channels.

It means that now the optogenetic control of the lysosome function through the regulation of pH over a wide range allows studying a wide range of fundamental lysosome depending cell functions.
